# Analysis of the effect of the scorpion toxin AaH-II on action potential generation in the axon initial segment

**DOI:** 10.1101/2023.10.06.561226

**Authors:** Fatima Abbas, Laila Ananda Blömer, Hugo Millet, Jérôme Montnach, Michel De Waard, Marco Canepari

## Abstract

The toxin AaH-II, from the scorpion *Androctonus australis Hector* venom, is a 64 amino acid peptide that targets voltage-gated Na^+^ channels (VGNCs) and slows their inactivation. While at macroscopic cellular level AaH-II prolongs the action potential (AP), a functional analysis of the effect of the toxin in the axon initial segment (AIS), where VGNCs are highly expressed, was never performed so far. Here, we report an original analysis of the effect of AaH-II on the AP generation in the AIS of neocortical layer-5 pyramidal neurons from mouse brain slices. After determining that AaH-II does not discriminate between Na_v_1.2 and Na_v_1.6, i.e. between the two VGNC isoforms expressed in this neuron, we established that 7 nM was the smallest toxin concentration producing a minimal detectable deformation of the somatic AP after local delivery of the toxin. Using membrane potential imaging, we found that, at this minimal concentration, AaH-II substantially widened the AP in the AIS. Using ultrafast Na^+^ imaging, we found that local application of 7 nM AaH-II caused a large increase in the slower component of the Na^+^ influx in the AIS. Finally, using ultrafast Ca^2+^ imaging, we observed that 7 nM AaH-II produces a spurious slow Ca^2+^ influx via Ca^2+^-permeable VGNCs. Molecules targeting VGNCs, including peptides, are proposed as potential therapeutic tools. Thus, the present analysis in the AIS can be considered a general proof-of-principle on how high-resolution imaging techniques can disclose drug effects that cannot be observed when tested at the macroscopic level.

## Introduction

Since its isolation from the venom of the scorpion *Androctonus australis hector*, the alpha-toxin AaH-II was found to bind with very high affinity to rat brain axolemma [1]. Specifically, AaH-II targets diverse voltage-gated Na^+^ channels (VGNCs) [2] with nanomolar affinity [3], and it prolongs their inactivation in different systems [4,5] by trapping a deactivated state [6]. After the bite of the scorpion, the toxin causes paralysis, cardiac arrhythmia and death in mammals [7], but it may also exacerbate the systemic inflammatory response and promote the development of lung injury [8,9]. However, low doses of the toxin can generate an immunogenic response leading to expression of antibodies and this response can be used to create protective therapies against scorpion envenoming by designing constructs either with fusion proteins [10] or nanoparticles [11]. Beside the effects of toxicity, natural peptides targeting VGNCs have been proposed as the basis for understanding physiological processes or novel potential therapeutic strategies. For example, a similar scorpion toxin, Amm VIII, was reported to have hyperalgesic effects triggered by gain-of-function of VGNCs [12], while Hm1a, a selective activator of Na_v_1.1, the principal VGNC expressed in GABAergic interneurons, could rescue mouse models of epileptic Dravet syndrome from seizures and immature death [13]. Most of these effects are concentration dependent and while low concentrations may have therapeutic effects, high concentrations can be detrimental. Thus, given the interest in designing novel peptides targeting VGNCs and inspired by animal toxins, the fine characterisation of the effects of these peptides at minimal doses becomes important, in particular in the central nervous system.

In the brain, VGNCs are highly expressed in the axon initial segment (AIS) [14,15] where they trigger the generation of the action potential (AP) [16,17]. It is therefore expected that the effect of a VGNC activator is stronger in this neuronal compartment where the channel density is higher with respect to the other sites. In neocortical pyramidal neurons, the VGNC isoforms that mediate AP generation are Na_v_1.2 and Na_v_1.6 [18], which are distinguished by different biophysical properties [19]. Recently, using a cutting-edge technique to record fast Na^+^ currents in the AIS [20], in combination with ultrafast membrane potential (V_m_) imaging [21] and Ca^2+^ imaging [22,23] techniques, we were able to characterise the Na^+^ and Ca^2+^ currents mediated by Na_v_1.2 and the way in which its functional interaction with the BK Ca^2+^-activated K^+^ channel shapes AP generation in the AIS of mouse neocortical layer-5 (L5) pyramidal neurons [24]. Here, using the same approach in the same preparation, we characterised the effect of the mutated toxin AaH-IIR62K (AaH-II) on the generation of the AP at a concentration that produces the minimal perturbation of the somatic AP. We show that, in contrast of the VGNC inhibitor G^1^G^4^-huwentoxin-IV used in the previous study, AaH-II does not distinguish between Na_v_1.2 and Na_v_1.6. Yet, we found an effect that can be attributed to the exquisite permeability to Ca^2+^ of Na_v_1.2 [25].

## Results

The first set of experiments was a dose-response analysis of the effect of AaH-II in HEK293 cells, expressing either human Na_v_1.2 or human Na_v_1.6, using automated patch-clamp recordings, similarly to what it was done in previous studies [26,24]. This analysis was necessary to establish the efficacy of the toxin on the two VGNC isoforms expressed in L5-pyramidal neurons [18]. Na^+^ currents were induced in voltage clamp by 50-ms pulses from -100 mV to 0 mV. The examples reported in Fig.1A show that bath perfusion of AaH-II, at the concentrations of 0.3 nM or 10 nM, prolonged the duration of both Na_v_1.2-mediated or Na_v_1.6-mediated Na^+^ currents by delaying their inactivation. The test was performed in distinct groups of cells at concentrations ranging from 0.01 nM to 10 nM. To quantify the effect of the toxin, we calculated the half inactivation time (T50) defined as the time at which the Na^+^ current reaches 50% of the peak current (see red arrows in Fig.1A). As shown in the plots of Fig.1B, AaH-II targeted both Na_v_1.2 and Na_v_1.6 VGNCs, poorly discriminating between them, and it produced a consistent effect on both channels in the concentration range 0.3-10 nM. The effect of the toxin was already maximal at 3 nM concentration.

**Figure 1.**
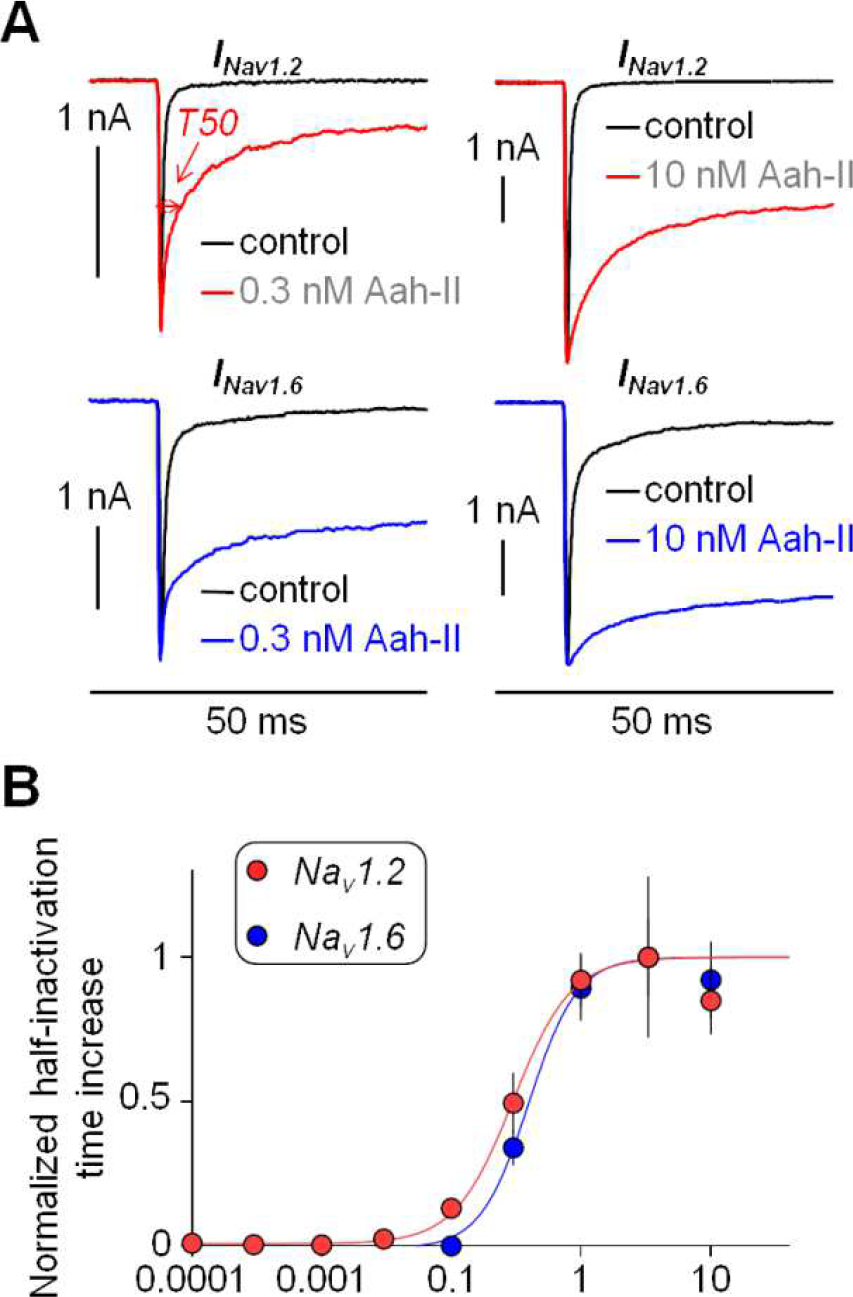
Dose-response analysis of the effect of AaH-II on Na_v_1.2 and Na_v_1.6 in HEK293 cells expressing either one or the other VGNC. (**A**) Na_v_1.2 or Na_v_1.6 currents in control solution and after addition of 0.3 nM or 10 nM of AaH-II. Red and blue traces are the Na_v_1.2 and Na_v_1.6 currents, respectively, in the presence of the toxin; on the top-left panel the current parameter T50 with 0.3 nM AaH-II is indicated with the red arrows. (**B**) Mean ± SEM of the normalized half inactivation time increase following AaH-II addition at different concentrations for Na_v_1.2 and for Na_v_1.6 currents. Values reported in the plot were as follows. For Na_v_1.2 (red lines and symbols): 0.01 nM – 0.004 ± 0.006 (N=7); 0.03 nM – 0.02 ± 0.01 (N=6); 0.1 nM 0.13 ± 0.04 (N=4); 0.3 nM 0.49 ± 0,11 (N=5); 1 nM – 0.92 ± 0.09 (N=9); 3 nM – 1 ± 0.28 (N=4); 10 nM – 0.85 ± 0.12 (N=6). For Na_v_1.6 (blue lines and symbols): 0.01 nM – 0.01 ± 0.005 (N=7); 0.03 nM – 0.007 ± 0.005 (N=9); 0.1 nM -0.0009 ± 0.006 (N=2); 0.3 nM – 0.34 ± 0.06 (N=20); 1 nM – 0.89 ± 0.11 (N=16); 3 nM – 1 ± 0.13 (N=20); 10 nM – 0.92 ± 0.13 (N=17). AaH-II does not discriminate between the two VGNC isoforms.

With this acquired information, we next aimed at establishing the effective concentrations changing the shape of the AP of L5 pyramidal neurons in brain slices, recorded by using the patch clamp technique in current clamp mode. Specifically, we elicited APs by setting the initial V_m_ to -80 mV and by injecting 2-3 nA current pulses of 3-4 ms duration, using a local delivery approach described in our previous report to apply the toxin [24]. Since in this study the equivalent concentration producing an effect in the brain slice was ∼40 times larger with respect to the experiments in HEK293 cells, we initially tested the effect of local delivery of 40 nM AaH-II on the somatic AP of L5 pyramidal neurons, expecting at this concentration a maximal effect. As shown in the example of Fig.2A, AaH-II delivery at 40 nM distorted dramatically the AP kinetics by increasing its amplitude and by widening its shape. The same qualitative effect was observed in all 5 cells tested with this concentration. Thus, in the example of Fig.2A, AaH-II was tested at three lower concentrations (5 nM, 10 nM and 20 nM) that produced a progressively larger effects evaluated by the percent change in AP amplitude and by the percent change in AP width, quantified by the time at which V_m_ > -30 mV during the AP. The same assessment was repeated in N = 8 cells (Fig.2C). Whereas the change in amplitude was not significant at any concentration, the change in width was 12.83 ± 7.77% and 25.41 ± 12.75% for the AaH-II delivery at 10 nM and 20 nM respectively, in both cases statistically significant (p < 0.01, paired t-test). At 5 nM, a detectable change in the AP width was observed only in 50% of the cells.

**Figure 2.**
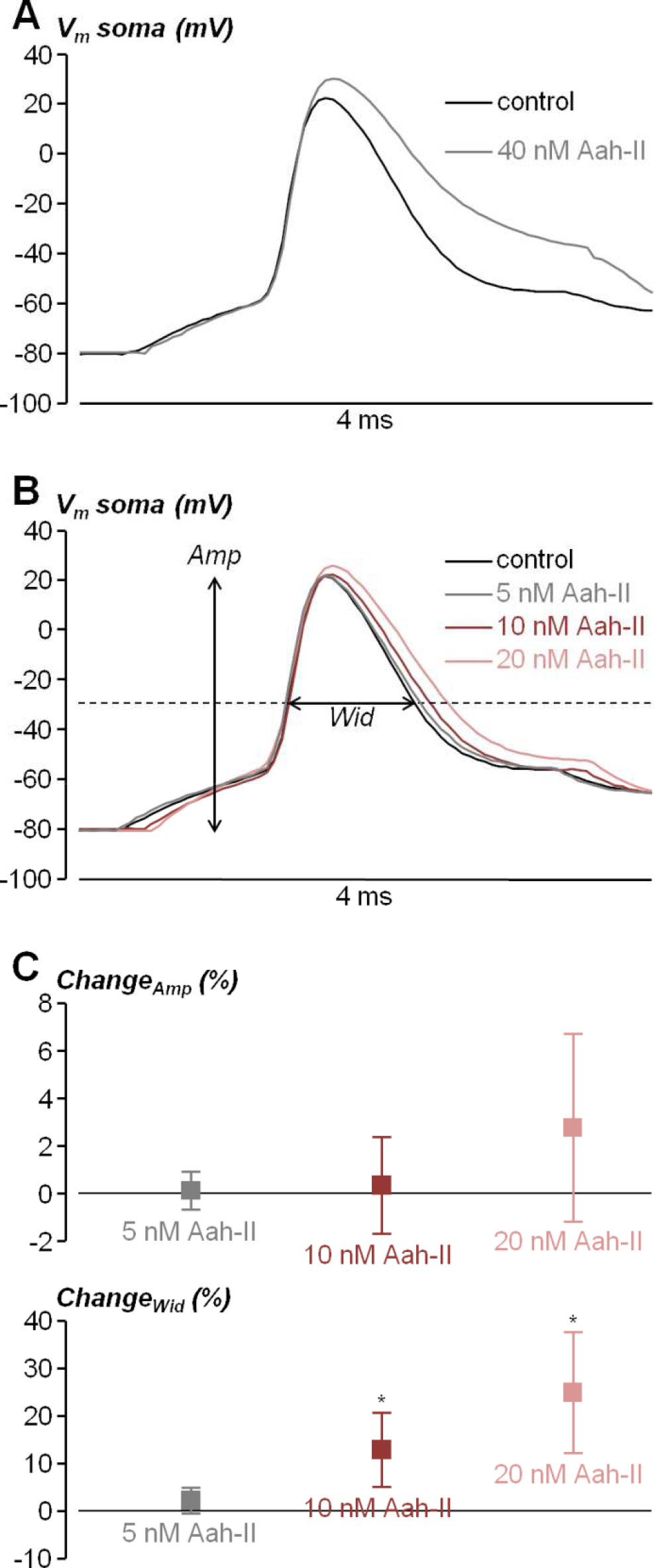
Dose-response analysis of the effect of AaH-II on the somatic AP from L5 pyramidal neurons in brain slices. (**A**) Somatic AP elicited by current pulse injection in a L5 pyramidal neuron in control condition (black trace) and after local delivery of 40 nM AaH-II (grey trace). (**B**) Somatic AP elicited by current pulse injection in another L5 pyramidal neuron in control condition (black trace) and after local delivery of 5 nM (grey trace), 10 nM (red trace) and 20 nM (pink trace) AaH-II. The amplitude (*Amp*) and width (*Wid*), used for the analysis, are indicated for the control trace. (**C**) Mean ± SD (N=8 cells) of the *Amp* and *Wid* changes following AaH-II addition at 5 nM (grey lines and symbols), 10 nM (red lines and symbols) and 20 nM (pink lines and symbols). Values reported in the plot were the followings. For the *Amp* parameter: 5 nM - 0.13±0.81%; 10 nM - 0.33±2.02%; 20 nM 2.80±3.94%. For the *Wid* parameter: 5 nM - 2.34±2.65%; 10 nM - 12.83±7.77%; 20 nM 25.41±12.75%. “^*^” indicates that the change in the *Wid* parameter was significant at 10 nM (p = 0.0094, paired t-test) and at 20 nM (p = 0.0034, paired t-test).

We therefore performed the next experiments by delivering AaH-II at the intermediate concentration of 7 nM in cells filled with the voltage-sensitive dyes JPW1114, in order to simultaneously record the somatic AP with the patch clamp electrode and the AP in the AIS optically. To standardise the analysis and minimise the variability along the AIS, we averaged fluorescence over a 20 µm segment between ∼10 µm and ∼30 µm from the soma. As shown in the example of Fig.3A, whereas local delivery of 7 nM AaH-II produced an increase in somatic AP width of ∼7%, it produced in the same cell an increase in the corresponding axonal AP width of ∼40%. The combined analysis of the somatic and axonal AP following the delivery of 7 nM AaH-II was repeated in N = 8 cells (Fig.3B). In the soma, a significant change in the AP width (11.50 ± 6.81%) was measured. Thus, we could refer to 7 nM as the minimal concentration producing a consistent effect on the somatic AP after local delivery. In the AIS, the change in AP width was 37.24 ± 16.13%. But most importantly, the effect on the axonal AP was significantly larger than the effect on the somatic AP in the cells tested (p < 0.01, paired t-test).

**Figure 3.**
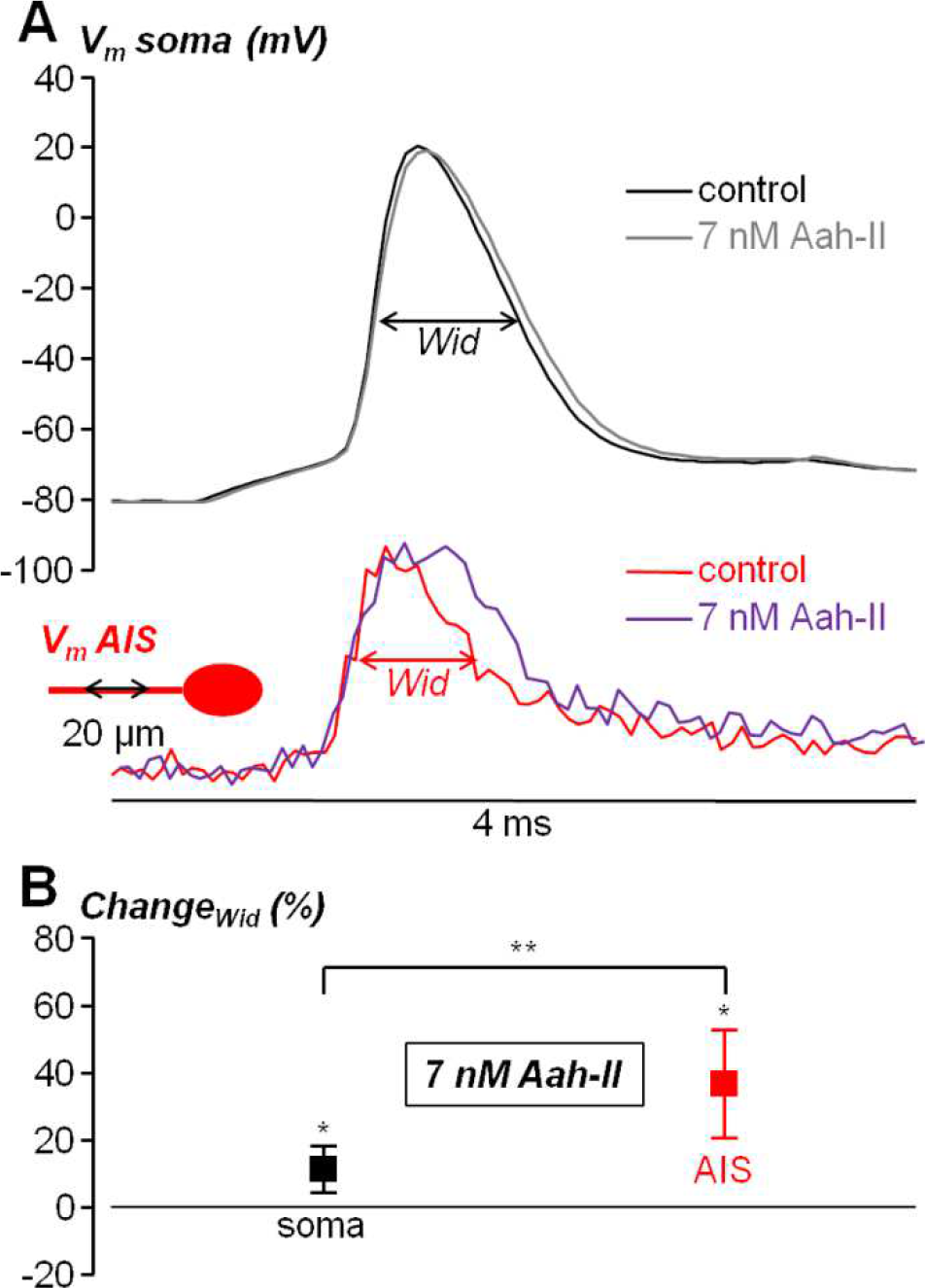
Effect of local delivery of 7 nM AaH-II on the somatic and AIS AP in L5 pyramidal neurons. (**A**) On the top, somatic AP elicited by current pulse injection in a L5 pyramidal neuron in control condition (black trace) and after local delivery of 7 nM AaH-II (grey trace). On the bottom, AIS (∼10-30 µm from the soma) AP in the same cell in control condition (red trace) and after AaH-II delivery (purple trace). *Wid* parameters used for the analysis are indicated for the soma and AIS relative to the control traces. (**B**) Mean ± SD (N=9 cells) of the *soma* (black lines and symbols) and AIS (red lines and symbols) *Wid* changes following AaH-II addition. Values reported in the plot were 11.50±6.81 for the soma and 37.24±16.13 for the AIS. “^*^” indicates that the change was significant for the soma (p = 0.0095, paired t-test) and for the AIS (p = 0.0034, paired t-test). “^**^” indicates that the change was significantly larger in the AIS (p = 3.4·10^-4^, paired t-test).

The result reported in Fig.3 could be attributed to the fact that in the AIS, where the AP is generated, the density of VGNCs is higher than in the rest of the cell [14]. This means that the analysis of the effect of the toxin on the AIS is capable of disclosing major effects that cannot be revealed by analysing the somatic AP. In keeping with the result reported in Fig.3B, we next assessed the effect of AaH-II delivery at minimal concentration on Na^+^ influx and Ca^2+^ influx in the AIS associated with the generating AP. In these two sets of experiments, cells were filled through the patch pipette either with the Na^+^ indicator ING-2, or with the Ca^2+^ indicator Oregon-Green BAPTA-5N (OG5N), and fluorescence transients associated with the AP, in the same AIS as in Fig.3, were recorded at 10k frames/s. As shown in the example of Fig.4A, AaH-II delivery caused a large increase in the late Na^+^ influx, corresponding to an increase in the non-inactivating Na^+^ current. This phenomenon was consistently observed in N = 9 cells where this test was performed. In the example of Fig.4B, AaH-II delivery caused instead a small increase in Ca^2+^ influx, corresponding to a widening of the Ca^2+^ current. Again, this phenomenon was consistently observed in N = 9 cells where this test was performed. We calculated mean ± SD of the change in the maximum fluorescence transients (Fig.4C) for Na^+^ (85.2 ± 25.9%) and Ca^2+^ (11.1 ± 5.3%) and we established that for both signals the increase produced by the toxin was significant (p < 0.01, paired t-test).

The fact that the change in Na^+^ influx was larger is not surprising since Ca^2+^ influx is mostly mediated by voltage-gated Ca^2+^ channels in the AIS [27]. Yet, we have previously demonstrated that the VGCC-component of the Ca^2+^ influx decreases with the distance from the soma and that the Na_v_1.2-mediated Ca^2+^ component is dominant in the distal part of the AIS [24]. Thus, we hypothesised that the change in the AP waveform had negligible effect on the VGCC-component of the Ca^2+^ influx and that the Ca^2+^ transient increase produced by AaH-II delivery was due to an increase in the Na_v_1.2-mediated Ca^2+^ component. To assess this hypothesis, we reanalysed the Ca^2+^ imaging dataset distinguishing from the distal AIS (30-40 µm from the soma) and proximal AIS (5-15 µm from the soma). In the example of Fig. 5A, AaH-II local delivery caused an increase in Ca^2+^ influx in the distal AIS, but not in the proximal AIS. As shown in Fig. 5B, in control condition, the peak of the distal Ca^2+^ current occurred before the peak of the proximal Ca^2+^ current. Then, the distal Ca^2+^ current (but not the proximal Ca^2+^ current) was widened in shape by the delivery of the toxin. In the 9 cells tested, the mean ± SD of the change in the maximum Ca^2+^ transients (Fig. 4C) was 18.7 ± 10.3% in the distal AIS and 0.0 ± 3.0% in the proximal AIS. A significant increase in Ca^2+^ transient was measured only in the distal AIS (p < 0.01, paired t-test). The spatial distribution of the Ca^2+^ transient increase can be also visually appreciated in Supplemental Movie S1. The additional Ca^2+^ signal produced by the delayed inactivation of Na_v_1.2 is slower than the control Ca^2+^ signal and it can therefore be considered a spurious component. Overall, the results presented in this study show how analysis of signalling in neuronal compartments of high VGNC density is necessary to unravel the effects of molecules targeting these channels, which are hidden in standard AP analysis from the cell bodies.

**Figure 4.**
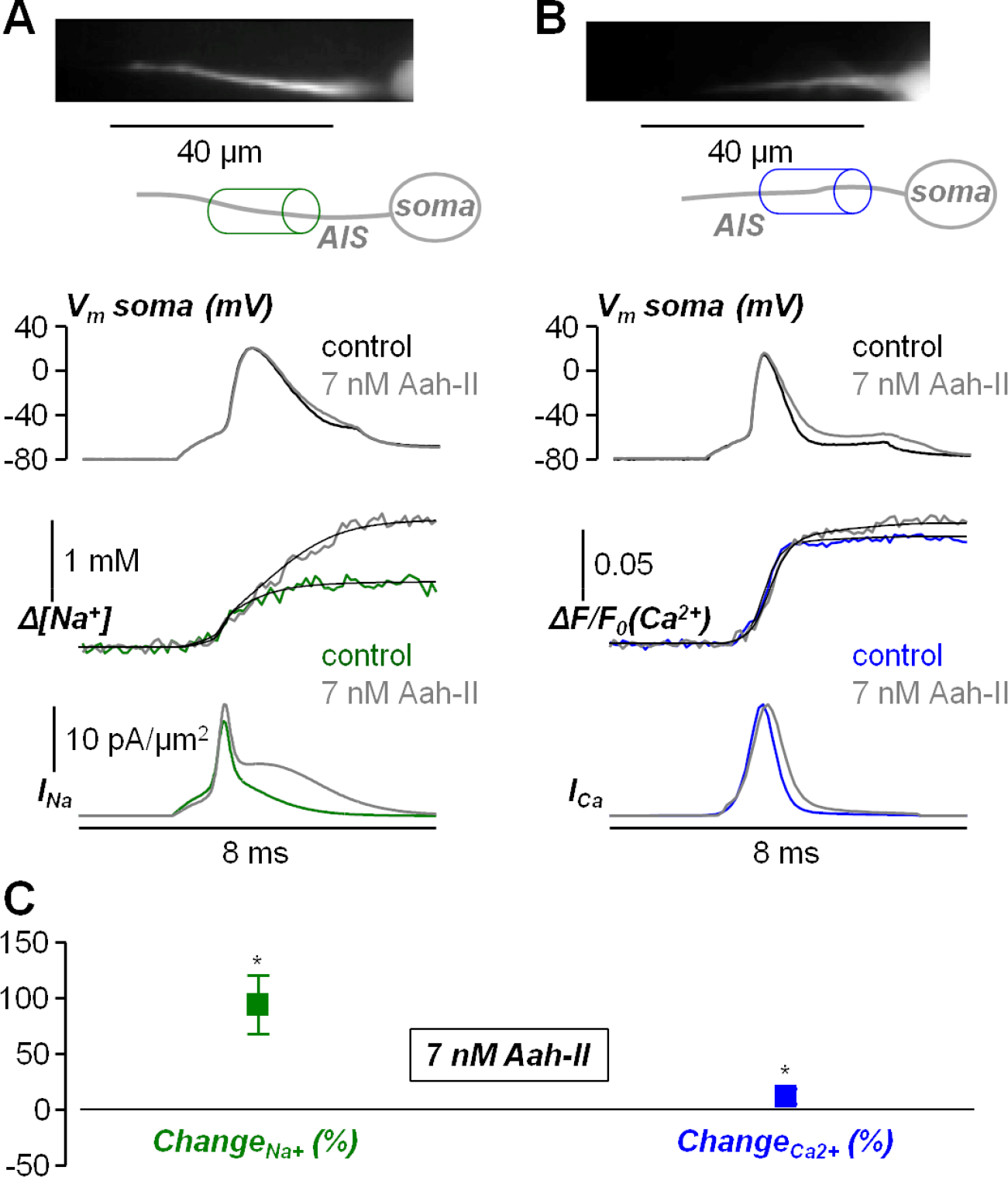
Effect of local delivery of 7 nM AaH-II on the Na^+^ and Ca^2+^ influx associated with the AP in the AIS. (**A**) Top, fluorescence image (Na^+^ indicator ING-2) of the AIS and its reconstruction with a region of interest indicated. Middle, somatic AP in control solution (black trace) and after delivering 7 nM of AaH-II (grey trace). Bottom, associated Na^+^ transients fitted with a model function to calculate the Na^+^ current (*I*_*Na*_) in control solution (green trace) and after delivering 7 nM of AaH-II (grey trace). (**B**) Top, fluorescence image (Ca^2+^ indicator OG5N) of the AIS and its reconstruction with a region of interest indicated. Middle, somatic AP in control solution (black trace) and after delivering 7 nM of AaH-II (grey trace). Bottom, associated Ca^2+^ transients fitted with a model function to calculate the Ca^2+^ current (*I*_*Ca*_) in control solution (blue trace) and after delivering 7 nM of AaH-II (grey trace). (**C**) Mean ± SD (N=9 cells) of the Na^+^ (green lines and symbols) and Ca^2+^ (blue lines and symbols) changes following AaH-II addition. Values reported in the plot were 85.2 ± 25.9%for Na^+^ and 11.1 ± 5.3% Ca^2+^. “^*^” indicates that the change was significant for the Na^+^ (p = 1.07·10^-6^, paired t-test) and for Ca^2+^ (p = 1.58·10^-5^, paired t-test).

**Figure 5.**
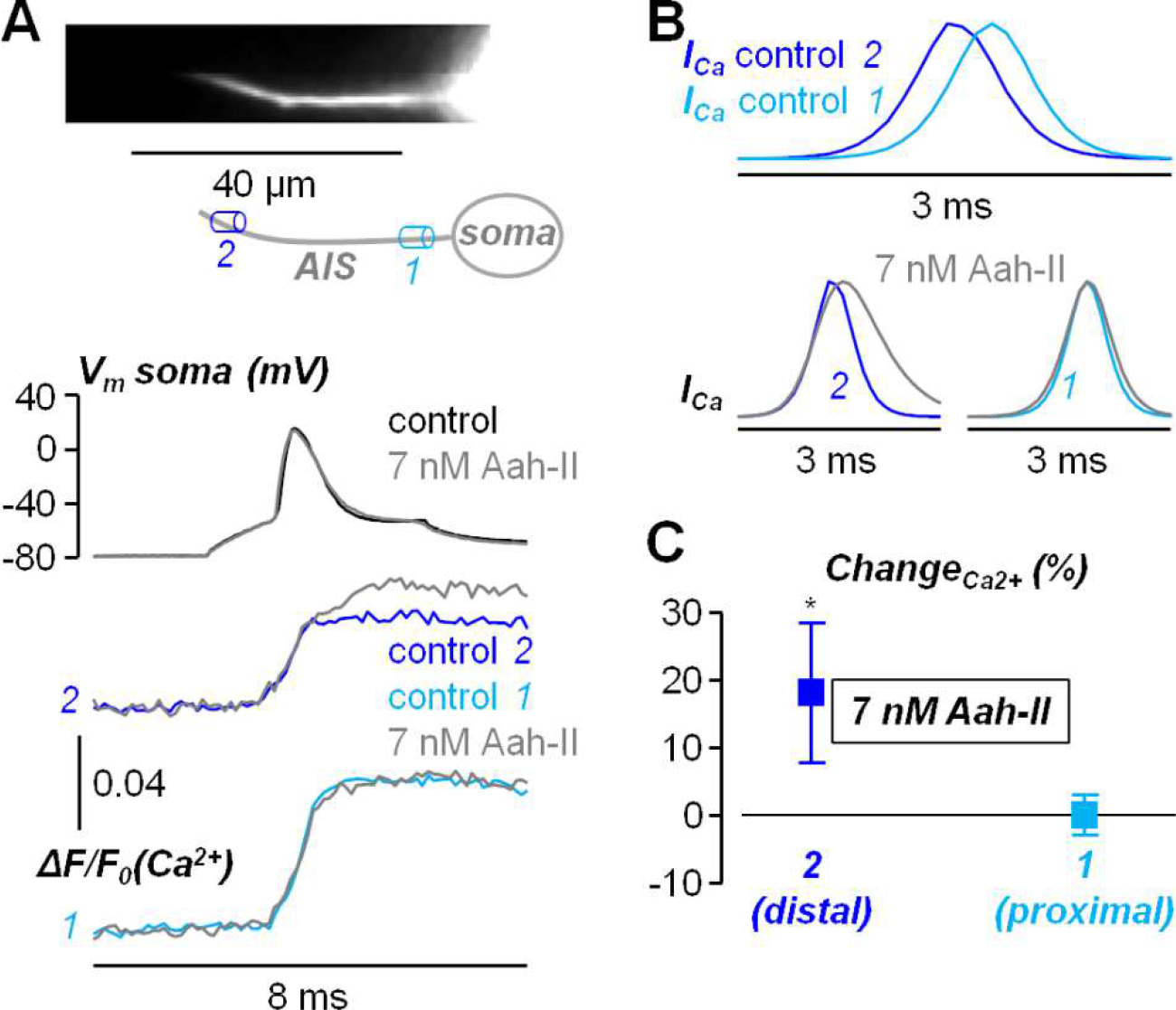
Spatial effect of AaH-II delivery on the Ca^2+^ influx associated with the AP in the AIS. (**A**) Top, Ca^2+^ fluorescence image of the AIS and its reconstruction with a proximal region (*1*) and a distal region (*2*) indicated. Middle, somatic AP in control solution (black trace) and after AaH-II delivery (grey trace). Bottom, associated Ca^2+^ transients in *2* and *1* in control solution (dark trace and light blue trace respectively) and after delivering 7 nM of AaH-II (grey traces). (**B**) Top, superimposed Ca^2+^ currents (*I*_*Ca*_) in *2* and *1* showing the different kinetics. Bottom, *I*_*Ca*_ in *2* and *1* in control solution (dark trace and light blue trace respectively) and after AaH-II delivery (grey traces). (**C**) Mean ± SD (N=9 cells) of the Ca^2+^ changes in distal AIS regions (30-40 µm from the soma, dark blue lines and symbols) and in proximal AIS regions (5-15 µm from the soma, light blue lines and symbols). Values reported in the plot were 18.7 ± 10.3% in the distal AIS and 0.0 ± 3.0% in the proximal AIS. “^*^” indicates that the change was significant in the distal AIS (p = 0.0011, paired t-test).

## Discussion

The first important finding that we report in this analysis is that local application of AaH-II produces a significant change in the shape of the somatic AP at a low 7 nM concentration, i.e. ∼10 times less with respect to the VGNC inhibitor G^1^G^4^-huwentoxin-IV used in a previous study [24]. This AaH-II effect corresponds to a slight widening of the somatic AP, but when we analysed the corresponding AP in the AIS, using a V_m_ indicator, the widening was significantly larger. Consistently with this result, local delivery of AaH-II caused a significant increase in Na^+^ influx associated with firing activity, that can be recorded in the AIS [28,29]. Not surprisingly, since the primary effect of AaH-II was to prolong VGNC inactivation, the larger influx could be attributed to the slow (or persistent) component of the Na^+^ current [30], which is normally larger for Na_v_1.6 in control condition [31]. When analysing Ca^2+^ influx associated with the AP, we found a significant increase after local delivery of AaH-II, that was however substantially smaller with respect to the increase in Na^+^ influx. It is known that a large component of Ca^2+^ influx in the AIS of L5 pyramidal neurons is mediated by voltage-gated Ca^2+^ channels [27], but it is also known that Na_v_1.2 is permeable to Ca^2+^ [25]. Thus, we hypothesised that the additional Ca^2+^ in the presence of the toxin could be mediated by Na_v_1.2. We have previously demonstrated that the Na_v_1.2-mediated Ca^2+^ current and the VGCC-mediated Ca^2+^ current can be clearly distinguished by the different kinetics, the first one peaking during the AP rising phase and the second one peaking in the AP falling phase [24]. In addition, we have demonstrated that the Na_v_1.2-mediated Ca^2+^ current is dominant in the distal part of the AIS where the VGCC-mediated Ca^2+^ current is weaker [24]. Consistently with this previous result and with the present hypothesis, we found that the increase in Ca^2+^ influx caused by AaH-II delivery occurred primarily in the distal part of the AIS. Yet, whereas in control condition the Na_v_1.2-mediated Ca^2+^ current is sharp and occurs during the rising phase of the generating AP [24], the wider Ca^2+^ current in the presence of AaH-II produced a late abnormal Ca^2+^ transient that can perturb axonal homeostasis.

From the methodological point of view, the present results indicate that the effect of a molecule targeting VGNCs should be analysed at the sites where the channels are highly expressed because substantial changes of physiological signals can be hidden when the analysis is performed at the full cellular scale. In the case of neurons, these sites are the AIS and the Ranvier nodes [32,33]. Mutations of VGNC genes are at the origin of a variety of channelopathies [34,35] making VGNC potential interesting targets for future therapies [36]. In addition, VGNCs are the potential targets of novel treatments for cancer [37] and inflammation-related diseases [38]. The present study represents a proof-of-principle on how our high-resolution imaging approach to explore AP generation in the AIS can be used to obtain relevant information on therapeutic candidates targeting VGNCs.

## Methods

### Ethical Approval

Experiments in brain slices were performed at the Laboratory of Interdisciplinary Physics in Grenoble in accordance with European Directives 2010/63/UE on the care, welfare and treatment of animals. Procedures were reviewed by the ethics committee affiliated to the animal facility of the university (D3842110001).

### Experiments in HEK293 cells expressing either Na_v_1.2 or Na_v_1.6

The mutated toxin AaH-IIR62K was the one described in a previous report [26] and it was produced by Smartox Biotechnology (Saint Egrève, France). Experiments on HEK293 cells were performed on stable lines expressing either human Na_V_1.2 or human Na_V_1.6 channels. Automated patch-clamp recordings in whole-cell configuration were performed using the SyncroPatch 384PE from Nanion (München, Germany) as previously described [26]. The intracellular solution contained (in mM): 10 CsCl, 110 CsF, 10 NaCl, 10 EGTA and 10 HEPES (pH 7.2, osmolarity 280 mOsm). The extracellular solution contained (in mM): 140 NaCl, 4 KCl, 2 CaCl_2_, 1 MgCl_2_, 5 glucose and 10 HEPES (pH 7.4, osmolarity 298 mOsm). Voltage clamp experiments were performed at a holding potential of −100 mV at room temperature (18-22°C) and currents were sampled at 20 kHz. To quantitatively assess the effect of the toxin, we calculated the parameter T50 defined as the time of 50% of inactivation between the peak of the sodium current and the residual current at the end of the depolarisation. For each cell, T50 was obtained after 15 min of AaH-II incubation and the change in the T50 parameter was obtained from the value prior to AaH-II incubation.

### Brain slices

The extracellular solution that we used contained (in mM): 125 NaCl, 26 NaHCO_3_, 1 MgSO_4_, 3 KCl, 1 NaH_2_PO_4_, 2 CaCl_2_ and 20 glucose, bubbled with 95% O_2_ and 5% CO_2_. Mice (C57BL/6j, 21-35 postnatal days old) were fed *ad libitum* until euthanasia. Animals were anesthetised by isoflurane inhalation and decapitated and the entire brain was quickly removed. Brain slices including the somato-sensory cortex (350 µm thick) were prepared using the procedure described in previous report [20,24], using a Leica VT1200 vibratome (Wetzlar, Germany). Slices were incubated at 37°C for 45 minutes and maintained at room temperature before use.

### Electrophysiology, imaging and drug delivery in brain slices

Slices were transferred to the recording chamber and patch-clamp recordings were made at the temperature of 32-34°C using a Multiclamp 700A (Molecular Devices, Sannyvale, CA). The intracellular solution contained (in mM): 125 KMeSO_4_, 5 KCl, 8 MgSO_4_, 5 Na_2_-ATP, 0.3 Tris-GTP, 12 Tris-Phosphocreatine, 20 HEPES, adjusted to pH 7.35 with KOH. For V_m_ recordings in the AIS, neuronal membranes were loaded with the voltage-sensitive dye D-2-ANEPEQ (JPW1114, 0.2 mg/mL, Thermo Fisher Scientific) for 30 minutes using a first patch clamp recording and then re-patched a second time with dye free solution. For Na^+^ or Ca^2+^ recordings in the AIS, either the Na^+^ indicator ING-2 (IonBiosciences, San Marcos, TX) or the Ca^2+^ indicator Oregon Green BAPTA-5N (Thermo Fisher Scientific) were added to the intracellular solution at the concentrations of 0.5 mM or 2 mM respectively. Somatic APs were elicited in current clamp mode by 2-3 ms current pulses of 2-3 nA through the patch pipette and electrical signals were acquired at 20 kHz. The bridge was corrected offline by using the recorded injected current and the measured V_m_ was corrected for -11 mV junction potential. V_m_ imaging experiments were performed using an imaging system described in recent reports [39,40], which was based on an Olympus BX51 microscope equipped with a 60X Nikon water immersion objective (NA = 1). V_m_ was excited using the 528 nm line of an LDI-7 laser (89 North, Williston, VT), band-pass filtered at 531 ± 40 nm and directed to the preparation using a 575 nm long-pass dichroic mirror. Fluorescence emission was long-pass filtered at >610 nm and acquired with a NeuroCCD camera (Redshirt Imaging, Decatur, GA) at 20 kHz. Na^+^ and Ca^2+^ imaging experiments were performed using an imaging system described in recent reports [20,24]. This system was based on an upright Scientifica SliceScope microscope equipped with a motorised XY translation stage and PatchStar manipulators (Scientifica, Uckfield, UK), and a 60X Olympus water immersion objective (NA = 1). Na^+^ or Ca^2+^ indicators were excited using the 520 nm line or the 470 nm line of a LaserBank (Cairn Research, Faversham, UK), band-pass filtered at 517 ± 10 nm or at 469 ± 17 nm and directed to the preparation using a 538 nm or a 495 nm long-pass dichroic mirror. Na^+^ or Ca^2+^ fluorescence was filtered at 559 ± 17 nm or at 525 ± 25 nm, demagnified by 0.5X and acquired at 10 frames/s by a DaVinci 2K CMOS camera (SciMeasure, Decatur, GA). To locally deliver AaH-II at a desired concentration in the proximity of the soma and AIS, we used a procedure described in a previous report [24]. Briefly, a pipette of 2-4 µm diameter was used to locally deliver AaH-II for 1 minute, by gentle pressure application. The effect of the toxin delivery was stable for at least 10 minutes after application, allowing the reported measurements.

### Data analysis

Data, from averages of 4-7 trials with identical somatic response, were analysed using custom-written code written in MATLAB. All optical recordings were corrected for photo-bleaching using multi-exponential fits of fluorescence transients in single trials without signal. The fractional change of fluorescence from the initial image (ΔF/F_0_) was calculated as previously described [24] and Na^+^ imaging experiments only ΔF/F_0_ was calibrated into a Na^+^ transient as described in a previous report [20]. ΔF/F_0_ transients were fitted with model functions and Na^+^ and Ca^2+^ currents estimated by calculating the time-derivative of the fit. The currents were not however considered in the quantitative analysis of the effects of the toxin.

### Statistics

The effects of AaH-II were quantified by calculating the percent change in the parameter analysed. To assess the consistency of an effect on groups of 8 or 9 cells, we performed a paired t-test and we considered 0.01 as the threshold to describe an effect as significant.

## Acknowledgements

This work was supported by the *Agence Nationale de la Recherche* through three grants (ANR-21-CE18-0042 – Nav12RESCUE; Labex *Ion Channels Science and Therapeutics:* program number ANR-11-LABX-0015 and National Infrastructure France Life Imaging “Noeud Grenoblois”) and by the *Fédération pour la Recherche sur le Cerveau* (FRC – Grant *Espoir en tête*, Rotary France). We are indebted to Fondation Leducq and the FEDER resources for financing the automated patch-clamp setup.

